# The African swine fever modelling challenge: objectives, model description and synthetic data generation

**DOI:** 10.1101/2021.12.20.473417

**Authors:** Sébastien Picault, Timothée Vergne, Matthieu Mancini, Servane Bareille, Pauline Ezanno

## Abstract

African swine fever (ASF) is an emerging disease currently spreading at the interface between wild boar and pig farms in Europe and Asia. Current disease control regulations, which involve massive culling with significant economic and animal welfare costs, need to be improved. Modelling enables relevant control measures to be explored, but conducting the exercise during an epidemic is extremely difficult. Modelling challenges enhance modellers’ ability to timely advice policy makers, improve their readiness when facing emerging threats, and promote international collaborations. The ASF-Challenge, which ran between August 2020 and January 2021, was the first modelling challenge in animal health. In this paper, we describe the objectives and rules of the challenge. We then demonstrate the mechanistic multi-host model that was used to mimic as accurately as possible an ASF-like epidemic, provide a detailed explanation of the surveillance and intervention strategies that generated the synthetic data, and describe the different management strategies that were assessed by the competing modelling teams. We then outline the different technical steps of the challenge as well as its environment. Finally, we synthesize the lessons we learnt along the way to guide future modelling challenges in animal health.

**Highlights:**

- The 1^st^ modelling challenge in animal health mimics ASF spread with synthetic data
- A mechanistic spatially-explicit stochastic model was developed to represent ASF spread and control
- Synthetic data concerned date and location of detected cases in pigs and wild boar
- Modelling ASF at the wildlife-livestock interface illustrates the reservoir role of wild fauna

## 1. Introduction

To sustainably raise livestock while respecting animal welfare, one must manage animal health, including infectious diseases which spread extensively between animal populations (Ezanno et al. 2020). Livestock diseases are currently being affected by changes at both the farm and global levels. Societal demand for more outdoor livestock farming may increase contacts between livestock and wildlife, and thus pathogen spread at this interface (Gortázar et al. 2007). Movements of persons, animals and animal products also are intensifying, and together with climate changes, are favouring the emergence and spread of new pathogens (Parham et al., 2015; Saker et al., 2004). These developments are impacting animal health and may affect public health when zoonoses are concerned.

African swine fever (ASF) is a good example of an emerging animal disease spreading at the interface between wildlife and livestock (Sánchez-Cordón et al., 2019; EFSA, 2021). This non-zoonotic viral disease originates from sub-Saharan Africa, where it is asymptomatically transmitted among warthogs and bushpigs. Wild boar and domestic pigs are susceptible hosts, with mortality rates of almost 100% for virulent strains such as the one currently circulating on the Eurasian continent (Dixon et al., 2020). Due to this high mortality rate, ASF has a tremendous impact on swine production, on the economy of livestock systems and on international trade, and neither a vaccine nor a treatment is available (Dixon et al., 2020). Therefore, since 21 April 2021, ASF has been categorized as an A+D+E disease under EU legislation. This means that immediate eradication measures must be taken as soon as the disease is detected in an EU member state (A), that the emergence of the disease in a country induces strong trade restrictions with the other member states (D), and that surveillance protocols are mandatory in all member states (E). In 2007, a highly virulent strain was introduced into Georgia (Rowlands et al. 2008) and then spread to the Russian Federation, from where it disseminated towards Europe and Asia (Dixon et al., 2020). It is now considered endemic in several countries of Eastern Europe and East and Southeast Asia. As the virus can spread internationally via geographical proximity or due to the movements of persons, swine and swine products, ASF has become one of the most important livestock infectious disease threat for most countries (Vergne et al., 2017).

Every health crisis, e.g., the recent pandemic of COVID-19 in humans (Holmdahl and Buckee, 2020), the foot-and-mouth disease epidemics in livestock in the UK (Keeling, 2005) and the ash dieback epidemics in European ash trees (Coker et al., 2019), highlights the need for robust epidemiological knowledge and predictive tools to better cope with health uncertainty. Developing models that forecast disease spread is key to better understand epidemics (Grassly & Fraser, 2008). Models are not new in epidemiology (Hamer, 1906), but with the computer revolution, and in the age of big data where information is shared almost in real time, this field has been revolutionized (Rosenfeld et al., 2013).

Models are highly valuable tools not only to anticipate what can happen under current conditions, but also to explore alternative scenarios regarding underlying assumptions, to anticipate environmental, ecological, and social changes, and to assess the effectiveness of control strategies. During new outbreaks, models are also useful for estimating unobservable parameters based on the first data records, as highlighted early in the COVID-19 pandemic with a huge modelling effort dedicated to forecast virus spread and assess public policies (Bertozzi et al., 2020), but also to guide data collection (Metcalf et al., 2020). Besides, studying mathematical properties of models provided powerful insights on their intrinsic limitations, e.g., the impact of specific assumptions on the possibility to estimate epidemiological parameters (Park et al., 2020) or a quantification of the time horizon beyond which the sensitivity to parameter values prevented from producing reliable forecasts (Castro et al., 2020).

In the case of ASF, simulation models have been developed to assess stricter regulations in Europe (Halasa et al., 2016). Their results highlighted the difficulty to control ASF spread in wild boar based on existing regulations only, and promoted the use of alternative measures (Lange, 2015), such as barriers and intensive hunting in fenced areas, as used in Belgium (EFSA, 2020). However, deciding which measures to implement remains a challenge, especially once an epidemic has started and decisions need to be taken promptly. To enhance global preparedness for ASF epidemics, the livestock/wildlife interface must be explicitly considered, and a capacity to assess different and combined control measures must be developed (Hayes et al., 2021).

A wide variety of models can be considered to represent an epidemiological situation (Chretien et al., 2014; Nsoesie et al., 2014; Holmdahl and Buckee, 2020). They are generally categorized as phenomenological models (e.g., ARIMA – autoregressive integrated moving average models, generalized linear models, survival models) and mechanistic models (e.g., SIR-like models, multi-agent systems). While the advantages and disadvantages of each model type are generally known, it is difficult to decide *a priori* whether one type is systematically more suitable than another for addressing a given question. Comparing models is therefore essential, but the task is difficult. Models may not use the same data or cover the same periods of the epidemic. The geographical areas over which predictions are made also can differ. These limitations appeared during the Ebola epidemics in West Africa (Chretien et al., 2015), and during influenza epidemics (Chretien et al., 2014; Nsoesie et al., 2014). In addition, a lack of cooperation between modellers and public health decision-makers is often deplored, particularly due to differences in timelines (a few days for decision makers whereas modellers need more time, especially when no model was developed in peace time), and to a lack of trust or to misunderstandings on the added-value of models and data, which is hard to address without long-term interactions (Metcalf et al., 2015). Modellers also report difficulties in modelling the spread of a disease in real time, especially in the early stages, when very limited data is available and strong uncertainties in transmission mechanisms persist (Van Kerkhove & Ferguson, 2012). More preparation is clearly needed to face these challenges (Johansson et al., 2019), better compare the predictive effectiveness of different models, advance the field of epidemiological modelling, and assist public health decision making.

To improve the accuracy of model predictions and cooperation between actors, the scientific community has developed relatively short competitions known as modelling challenges. The first challenge, organized in 1994, was on protein structure prediction (Friedberg et al., 2015). In epidemiological modelling, the first challenge was organized in 2013 on seasonal influenza in the US, and has since been annually renewed (Reich et al., 2019; Viboud and Vespignani, 2019). Three other challenges also were organized on Ebola (Viboud et al., 2018), Chikungunya (Del Valle et al., 2018), and Dengue (Johansson et al., 2019). So far, however, no challenge has involved an animal disease. Modelling the transmission of such infectious diseases needs to address specific features, including the spatial distribution of farms and the difficulty of monitoring infectious diseases in wildlife.

This paper aims to introduce the ASF modelling challenge (ASF-Challenge). It presents the objective of the challenge, the stochastic, agent-based and spatialized epidemiological model developed to represent ASF spread at the interface between domestic pigs and wild boar, and the synthetic data produced using this model to mimic an ASF epidemic for the ASF-Challenge.

The performance of the different models developed by participating teams to the ASF-Challenge are presented in the last article from this special issue (Ezanno et al. submitted).

## 2. Methods

### 2.1 ASF-Challenge characteristics

The preparation for the ASF-Challenge started in July 2019. This preparation phase was used to define the ASF-Challenge objectives, build the “mother model” that generated the synthetic data used in the challenge, and clarify what was expected from the participating teams. The ASF-Challenge itself was initially planned to take place between March and July 2020, but it was postponed due to the COVID-19 pandemic. It was finally launched on 27 August 2020 and lasted until 13 January 2021. Once the ASF-Challenge itself had ended, a series of internal workshops were organized to obtain feedback from the challenge teams and present the different models that were used. This also provided an opportunity to synthesize the challenge outputs and reflect on the lessons learnt. These are presented in the last article from this special issue (Ezanno et al. submitted).

Data provided to the challenge teams were generated by a detailed agent-based model that was fed with population data (spatial distribution of the host populations, movements of live pigs, etc.) and parametrized with key parameters defining transmission processes and intervention strategies, as described in detail hereafter. For the ASF-Challenge, we aimed to generate ASF-like epidemic trajectories with the following characteristics: first outbreak detected in a domestic pig farm at the vicinity of a forest area and less than 200 days after the first infection, more than 250 infected wild boar at the time of the first detection, outbreaks reported in both domestic pig farms and wild boar over the course of the epidemic, apparent epidemic duration of more than 100 days, progressive diffusion towards a forest area with a high density of wild boar and an apparent successful control of the disease (i.e., less than 500 infected wild boar outside the fence when implemented, more than 250 infected wild boar 110 days after the first detection but less than 30 infected wild boar 230 days after first detection) at the end of the ASF-Challenge.

To ensure that the model represented a realistic setting of hosts and could reproduce an ASF-like epidemic with sufficient accuracy, we relied on a small group of French experts with knowledge on pig production, wild boar ecology and African swine fever epidemiology. These experts suggested relevant data sources and discussed model assumptions (population distributions, movement data, transmission processes, etc.).

### 2.2 Data used to feed the model

The simulated epidemic occurred on a hypothetical island that was created by merging two French administrative regions, namely Auvergne-Rhone-Alpes and Occitanie. All polygons and land use data were obtained from the DIVA-GIS website (https://www.diva-gis.org/) and all land use types were aggregated into three types: agricultural, forest and urban areas. The spatial locations of simulated individual wild boar and domestic pig farms were based on land use data.

To simulate wild boar distribution, we obtained the hunting bags at the department level from the *Office Français de la Biodiversité*, and assumed that a hunting season would reduce the wild boar population by half, so that the total wild boar population size was twice the hunting bag size at the level of the whole island. The location of each of the 500,366 wild boar individuals was randomly simulated, assuming that 18%, 80%, and 2% of them would be in agricultural, forest, and urban areas, respectively. Their geographical coordinates were assumed to represent the centre of their home range. Their spatial distribution was then summarized as department-level hunting bags to mimic the type of data that would be available to modellers in a real situation. To do so, we assumed again that hunting bag sizes represented half the size of the wild boar population in each department.

To simulate a regionalization and a spatially heterogeneous distribution of the 4,775 pig farms registered in the two French regions, we assumed that 1) 33% of farms would be in the Auvergne-Rhone-Alpes region while the remaining 67% would be in the Occitanie region, and 2) 85%, 10%, and 5% of pig farms would be in agricultural, forest, and urban areas, respectively. Under these specific constraints, we assigned random geographical coordinates to each of the farms. Specific farm characteristics were assigned to each premise: commercial or backyard; breeder, finisher, or breeder-finisher; whether pigs had access to an outdoor area; farm capacity; the regrouping of different farms into the same pig company (931 farms over 4,775 were randomly assigned to a company, resulting in 126 companies composed from 2 to 19 farms - 7.4 in average). All of this information was used to generate a biosecurity score to model variations in farm susceptibility to, and transmissibility of, ASF (see below).

Time-dependent live-pig trade movement data between farms were also simulated, assuming as in typical European intensive pig farming systems that breeder and breeder-finisher farms could send pigs to finisher or breeder-finisher farms; that a breeder-finisher farm was more likely to fatten its own pigs than to send them to another farm; that farms belonging to a pig company were more likely to send pigs to farms from the same company than to farms not belonging to the same company; and that the total number of pigs in a farm at a given time could not exceed the size of the farm.

### 2.3 Model used to produce synthetic epidemiological data

To produce these synthetic data, we needed a very detailed model that was expected to be more complex than any model used by the ASF-Challenge teams. Hence, we designed a stochastic mechanistic model which integrated the most up-to-date fine-grained knowledge and assumptions with regard to ASF spread and control in wild boar and domestic pigs. This model was in discrete time, with a time step of one day, spatially explicit and agent-based with three types of agents: the pig farm (as a compartment-based sub-model), the individual wild boar, and the whole hypothetical island as a metapopulation. The model was implemented using a recent modelling approach, EMULSION (Picault et al., 2019), which helps to make model components explicit and to reduce the amount of code required.

#### 2.3.1 Population dynamics and infection spread

The dynamics of the domestic pig population was determined only by the trade movements between farms depending on their type (breeders, finishers and breeder-finishers), neglecting natural mortality and replacing sold animals with the same number of new ones (representing births in breeders or breeder-finishers, and purchases in finishers or breeder-finishers), without considering the detailed farm structure explicitly (e.g., batch management). In wild boar, we considered natural mortality and hunting through constant rates. We considered no birth during the simulated period, as the epidemic was assumed to occur during the hunting season, i.e., after the reproductive period (Vetter et al 2020). The hunting rate was calculated to remove 50% of the wild boar population during the hunting season (in 8 months), consistent with how the hunting bags were used to generate the wild boar distribution (Jori et al., 2021). All of the parameters involved in the dynamics of pig farms and the wild boar population are provided in the supplementary information (Table S4).

All epidemiological units (pig farms and individual wild boar) were subject to the same infectious process, with the following states for animals in the units: susceptible (S), exposed (E) where animals started being infectious but were still asymptomatic, fully infectious and symptomatic (I). All infected animals eventually died, producing an infectious carcass (C). We also assumed that wild boar were subject to natural mortality, giving either a healthy (D) or infectious (C) carcass depending on their health state at death. The durations in states E, I, C, D were distributed exponentially. In pig farms, carcasses were removed the next day, whereas infectious (C) or healthy (D) wild boar carcasses could remain in the environment for several weeks or months until they naturally decomposed or were removed when found by a passer-by. Within pig farms, we assumed a frequency-dependent force of infection, exposed individuals contributing to half the level of infectious animals or infectious carcasses. We also assumed a higher transmission rate in backyard farms than in commercial farms, to account for the more compartmentalized contact structure induced by batch management in commercial farms. All of the parameters involved in the infection process are provided in supplementary information (Table S5).

For pig farms, we considered the following transmission pathways between epidemiological units: 1) arrival of an infected pig from an infected farm in another farm, through trade movements; 2) contact with an infectious wild boar; and 3) indirect contact with an infectious farm due to visits, exchange of agricultural material, etc. For wild boar, we considered: 4) contact with an infectious live wild boar; 5) contact with an infectious wild boar carcass; and 6) contact with an infectious pig farm. All of the transmission pathways assuming a contact with a farm were weighted by the biosecurity score of the farm.

Transmission based on trade relied on the pre-simulated animal movements, which determined the source and destination farms as well as the number of domestic pigs sold, with epidemiological states sampled randomly depending on their distribution in the source farm. All of the five other transmission pathways were spatially explicit, using exponential transmission kernel functions of the square of the distance between epidemiological units, considering different values for the kernel parameter (Table S1). We also assumed that contacts between wild boar and pig farms could occur only for farms providing outdoor access to pigs. Finally, we assumed that the contribution of infected pig farms to other susceptible epidemiological units was a function of their within-farm prevalence and their biosecurity level. The resulting forces of infection experienced by each possible epidemiological unit from other infectious units are summarized in the supplementary information (Table S2).

#### 2.3.2 Intervention strategies

The model represented several detection methods depending on the characteristics of each epidemiological unit. We assumed that all tests were perfectly sensitive and specific to focus on the interplay between intervention strategies, detection, and control. All of the parameters involved in the intervention strategies are provided in the supplementary information (SI3-5).

Prior to any primary case, detection relied only on passive surveillance. In wild boar, we assumed that each carcass could be found, reported and tested each day with a constant and low probability. In infected farms, each infected pig could be detected and tested each day while being in infectious state (I) with a constant probability, and at death. Detection probabilities varied depending on the farm type (higher in commercial farms, lower in backyard) and increased in all pig farms after the detection of the primary case to account for enhanced vigilance (Table S3).

The model incorporated all current European regulatory measures and triggered them as soon as a primary case was detected in the simulation. All animals in pig farms detected as infected were culled, and the delay between detection and culling was fixed. A protection zone and a surveillance zone were established (Table S6), both subject to a trade ban and increased vigilance (increased infection detection probabilities and biosecurity scores). Farms which had exchanged animals with a detected infected farm during the previous three weeks (called “traced farms” in what follows) were subjected to the same restrictions and vigilance as in the protection zone, with the same duration. Culled farms could be repopulated after a fixed period of time (Table S6).

Infected wild boar carcasses found in the environment were removed immediately and triggered an active search for other infected carcasses (Table S6). During this active search, each wild boar carcass (either infected or disease-free) could be found with an increased probability compared to carcasses outside the search zone. The detection of new infected carcasses prompted new active search operations (assuming no logistic constraints). After the detection of the primary case, it was also assumed that a fraction of hunted wild boar were tested (Table S6).

In addition to regulatory measures, alternative interventions were triggered in the simulation (Table S7). First, as the forest near the primary case could be considered a main threat for virus diffusion, 300 km of fences were installed 60 days after the detection of the primary case. The hunting pressure was increased within the fenced area as well as in a buffer area outside the fences, aiming to remove 90% of wild boar by the end of the hunting period (instead of 50% in other areas). In all areas with increased hunting, all hunted animals were tested, the active search for wild boar carcasses was suspended, and the probability of finding wild boar carcasses by passive surveillance was much higher than before. The increased hunting effort in the buffer area occurred for two months. Second, 90 days after the detection of the primary case and until the end of the simulation, when an infected wild boar was found (either through hunting or as a carcass), all animals from nearby farms (Table S7) were culled preventively and tested (leading to the installation of protection and surveillance zones and trade contact tracing if positive).

#### 2.3.3 Stochastic simulations, model outputs, and selected synthetic data

The virus was initially introduced shortly before the beginning of the hunting season (Table S4), through an exposed wild boar individual located near a forest close to the centre of the hypothetical island, all other epidemiological units being fully susceptible.

In addition to the numbers in each health state for each epidemiological unit, the model kept track of all events (infection; detection with date; cause of removal: active search, hunting, culling; test). However, the transmission trees were not recorded, for the sake of limiting the computational cost of the model. These fine-grained model outputs were used to (i) analyse the stochastic trajectories and select a relevant one for the challenge, and (ii) reconstruct simulated epidemiological data as time series given to the ASF-Challenge teams. We calculated the maximal geographical extent of the spatial distribution of cumulative cases, defined as a rectangle whose corners were the most extreme coordinates of the infected wild boar case (respectively pig farms) encountered during the challenge (increasing function). Finally, we assessed the probability of infection of each pixel (5×5 km^2^ squares), defined by the proportion of model repetitions where the pixel had at least one infected pig farm, live wild boar, or carcass.To find trajectories with ASF-like dynamics that were realistic enough for the challenge, 500 stochastic repetitions were run. First, simulations without any detection (40/500) or with a date of primary case detection later than 200 days after virus introduction (9/500) were discarded as unfit for the challenge. Among the 451 remaining simulations, five selection criteria were defined (SI6, Table S8). Seven replicates met all five selection criteria, among which the one actually selected for the ASF-Challenge was chosen at random.

## 3. Results

### 3.1 General predictions of the model

#### 3.1.1 Temporal dynamics

The infection dynamics was highly variable among the 451 stochastic model repetitions, despite similar initial conditions. This was especially true in wild boar where the number of cases typically ranged from 2,500 to 7,500 (Fig. 1). It grew rapidly in most of the repetitions, within a large range both for the date (circa 150 days) and amplitude (about 5,000 cases) of the epidemic peak. Regulated measures enabled the spread to be limited and to reach a plateau of cases before the fences were implemented. Fences and increased hunting were required to decrease the number of cases in wild boar and the associated exposure of pig farms. Finally, a rebound of the epidemics was highly probable after the increased hunting had stopped (while other control measures still were implemented).

**Figure 1.**
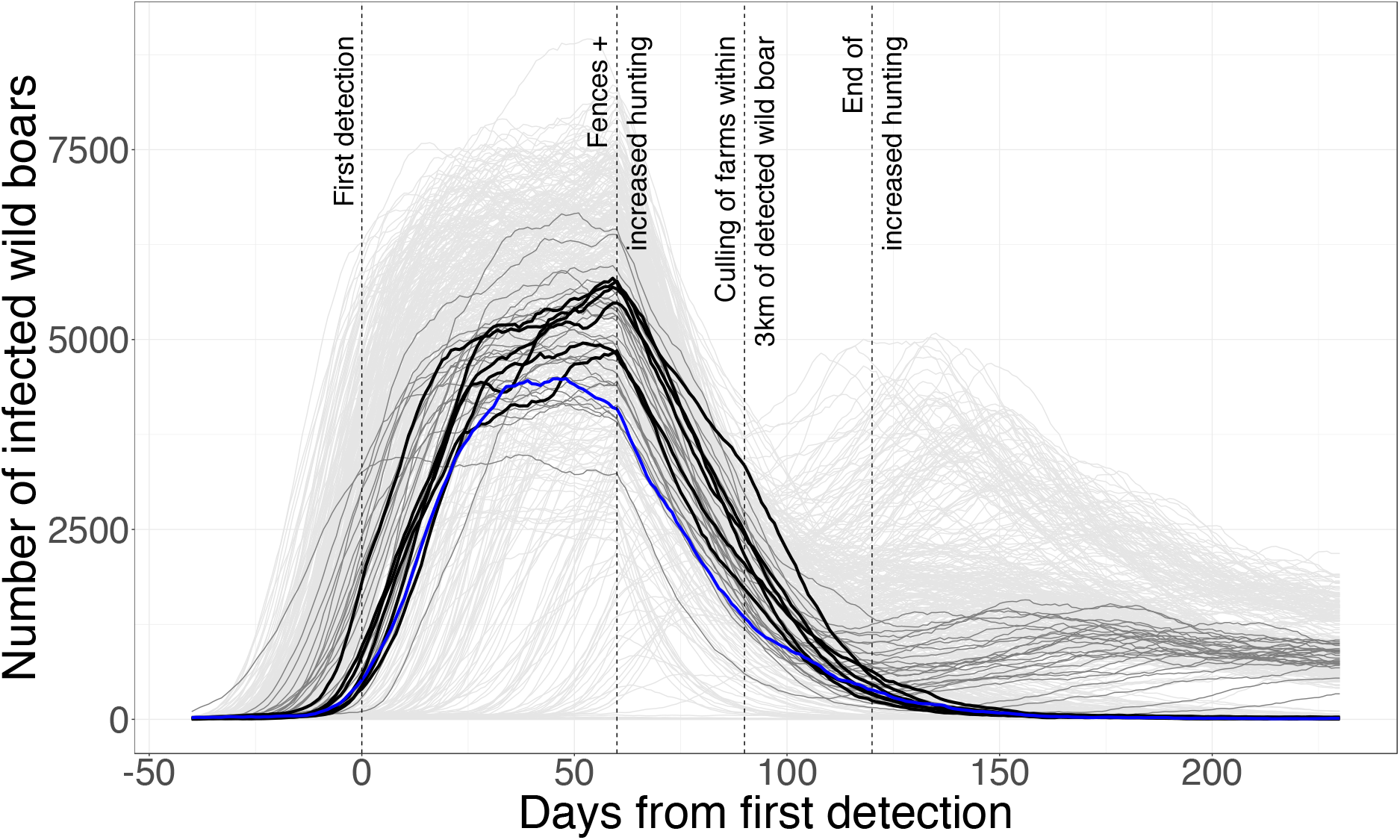
Temporal dynamics of the number of live infected wild boars (exposed + infectious) for 451 stochastic repetitions of the model. Blue: repetition selected for the ASF-Challenge; black: the 6 other repetitions meeting the 5 selection criteria; dark grey: the 34 repetitions meeting all the criteria except the last one (<30 infected wild boars 230 days after first detection). Vertical dotted lines: changes in control interventions.

The seven repetitions meeting all selection criteria (Tab. S8) showed small epidemics in pig farms (about 25 farms detected) and intermediate ones in wild boar about 4,000 cases – (Fig. 2). The last criteria (<30 infected live wild boar 230 days after first detection) led to what was almost disease fade-out (Fig. 1).

**Figure 2.**
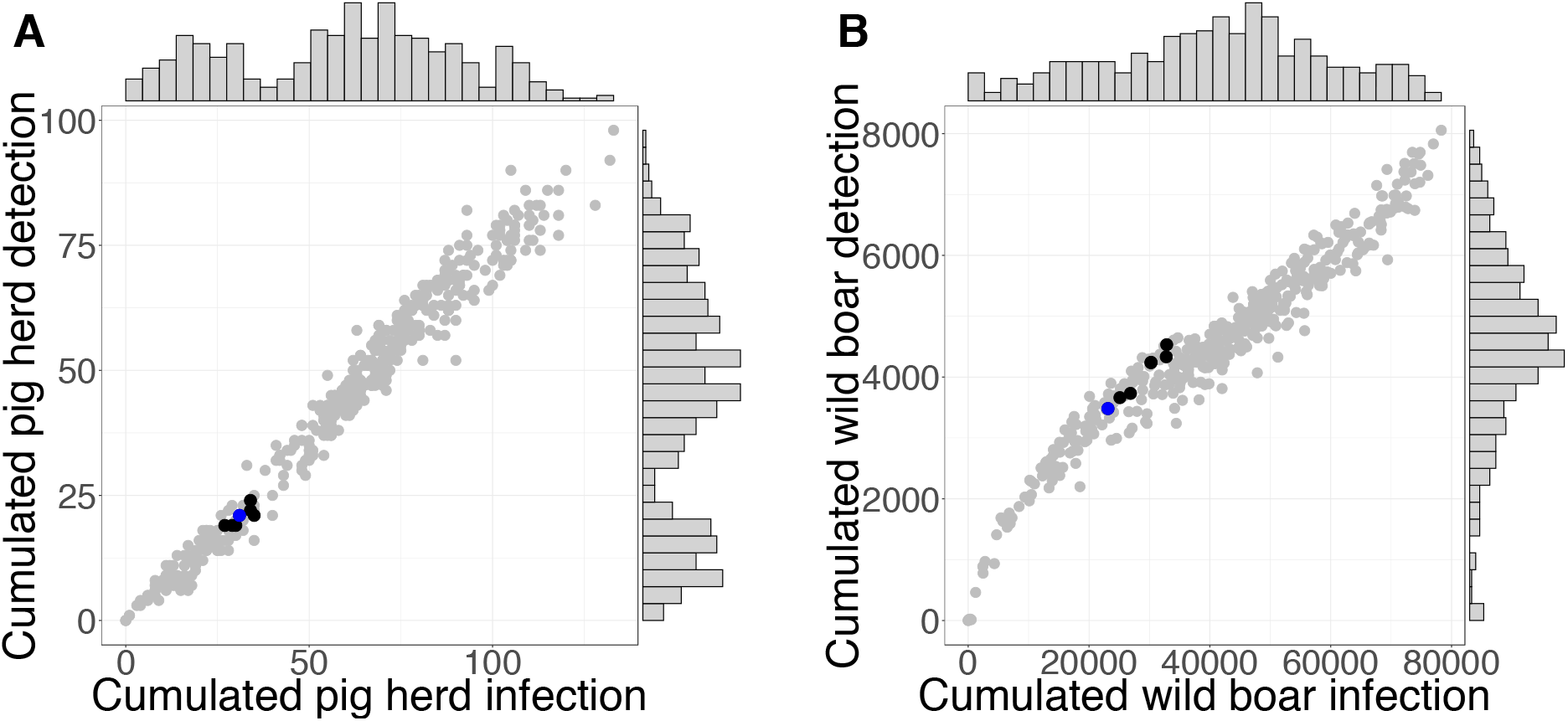
Distribution of the cumulative epidemic size and case detection up to 230 days after the first case detection for 451 stochastic repetitions of the model. A: pig farms; B: wild boar; blue: repetition selected for the ASF-Challenge; black: the 6 other repetitions meeting the five selection criteria; grey: other 444 repetitions.

#### 3.1.2 Spatial dynamics

The maximal geographical extent of the epidemics was highly variable among repetitions, both in the wild boar and pig populations (Fig. 3). The seven repetitions satisfying all selection criteria showed a low spatial spread in the wild boar population (Fig. 3B), very close to the surface of the fenced area. In contrast, they showed a much more variable spatial spread in pig farms (Fig. 3A), in relation with commercial movements of infected animals before source farms were detected. These long-distance spreading events in pig farms did not impact the spatial spread of the disease in wild boar, highlighting a low exposure of wild boar to infectious pig farms.

**Figure 3.**
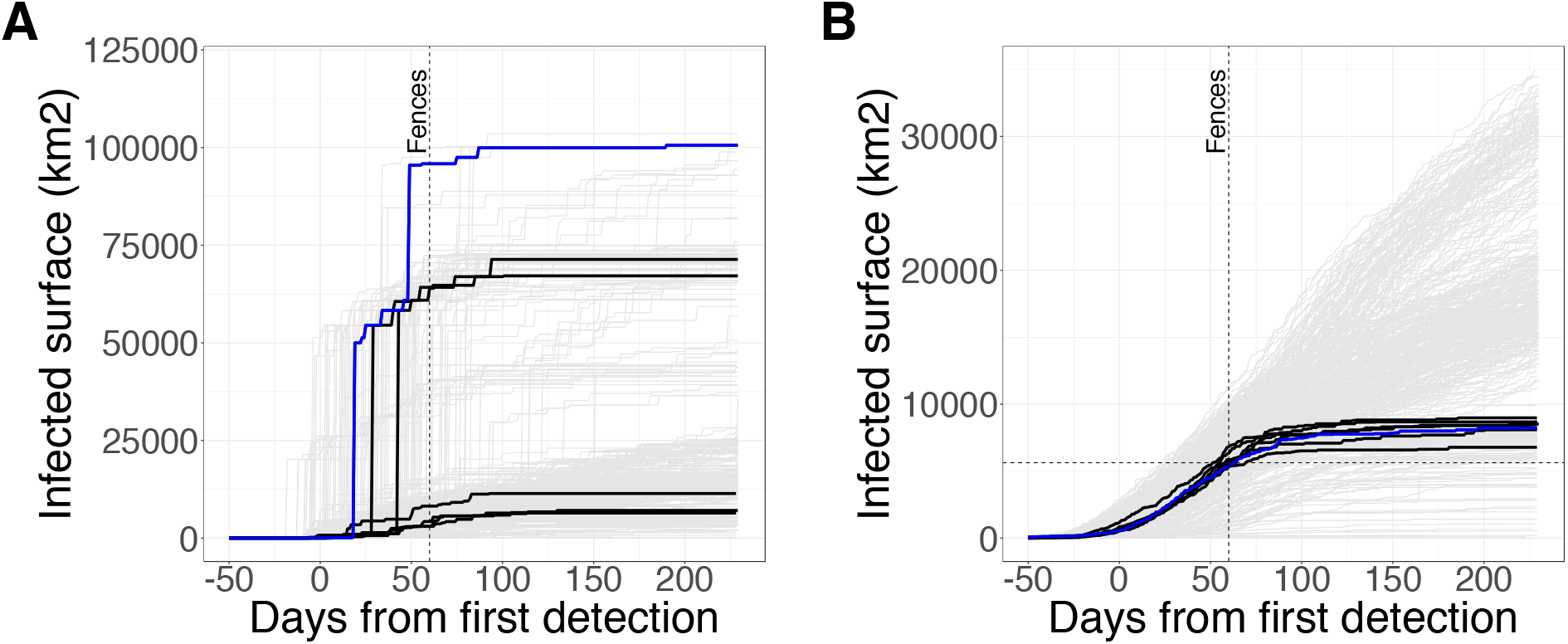
Temporal evolution of the maximal geographical extent of the spatial distribution of cumulative cases for 451 stochastic repetitions of the model. A: pig farms; B: wild boar; blue: repetition selected for the ASF-Challenge; black: the 6 other repetitions meeting the five selection criteria; grey: other 444 repetitions; vertical line: date of fences installation; horizontal dashed line: surface of the fenced area.

Despite a maximal geographical extent of the epidemics of nearly 100,000 km2, the cases were highly aggregated in the selected repetition, both for pig farms and wild boar individuals (Fig. 4). The local probability of infection was very high within the fenced area and on its western limit (where the primary case was introduced). It was high south of the fenced area, and low but not nil north and east of that area. It was nil everywhere else, except where a few pig farms has been infected in some of the stochastic model repetitions.

**Figure 4.**
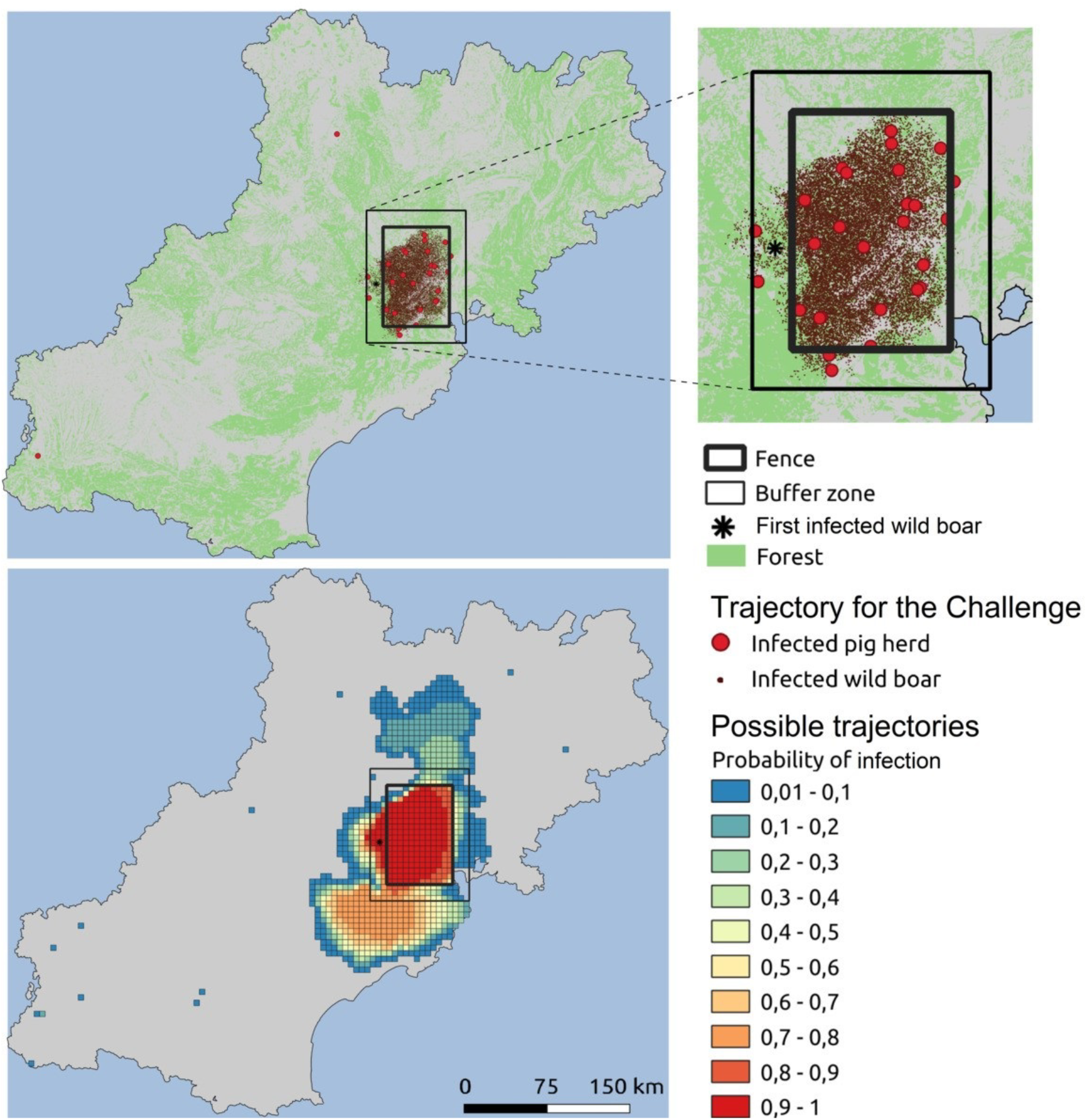
Spatial distribution of infected epidemiological units. Top: cumulated infected epidemiological units 230 days after the first case detection in the selected repetition (in red: pig farms, in brown: wild boar); Bottom: local probability of infection (i.e. probability that at least one domestic pig farm or one wild boar becomes infected over the course of the epidemic) calculated on the 451 stochastic repetitions of the model (trajectories presented on Figure 1), at a 5×5 km^2^ scale.

### 3.2 Data for players

Before the ASF-Challenge started, the ASF-Challenge teams were provided with the available population data, i.e. the hunting bag size per department, as well as the farm database that comprised the location and characteristics of the pig farms (except for their biosecurity score). To allow the teams to become familiar with the data format and start developing their analytical pipelines before the actual start of the ASF-Challenge, dummy datasets were released to the participating teams four weeks before the start of the challenge. This data included the live-pig movement data and the ASF surveillance outputs for a phase of four weeks. This step was extremely important to motivate the teams and optimize their readiness.

The challenge was organized in three different epidemiological phases, phases 1, 2, and 3, which started when 50, 80, and 110 days had passed since the first case detection, respectively. At the beginning of each phase, the ASF-Challenge teams were provided with the surveillance outputs and a situation report.

Similar to the RAPIDD Ebola forecasting challenge (Ajelli et al., 2018), we documented the data format by preparing “Read Me” documents describing the structure of all of the different datasets and the interpretation of all of the different variables within each dataset (available in the code repository, see supplementary information SI1).

#### 3.2.1 Surveillance outputs

The epidemiological information that was released to the teams comprised two datasets in CSV format: the list of cases (detected infected pig farms and wild boar) and the list of hunted wild boar that tested negative. Both datasets had the same structure. They included the farm ID (NA if a wild boar), the host type (pig farm or wild boar), the geographical coordinates, the infection detection method, the date of suspicion, the date of confirmation and the date of culling (NA if a wild boar). The farm ID was the key to retrieve a farm’s characteristics from the farm database.

Overall, the synthetic data provided to the teams relied on a few detected cases in pig farms, spread over the three phases of the challenge, and a massive observed epidemic in the wild boar population. The number of detected cases correlated strongly with the number of actual cases (Fig. 2), but only represented a fraction of the epidemic, especially in wild boar, where only about 15% of infected animals were detected (Fig. 5). After the implementation of fencing and increased hunting in the fenced area, the number of detected wild boar cases was much greater. Before the implementation of fencing, the main ASF detection stream in wild boar was the active search of carcasses, while it became the testing of hunted animals after.

**Figure 5.**
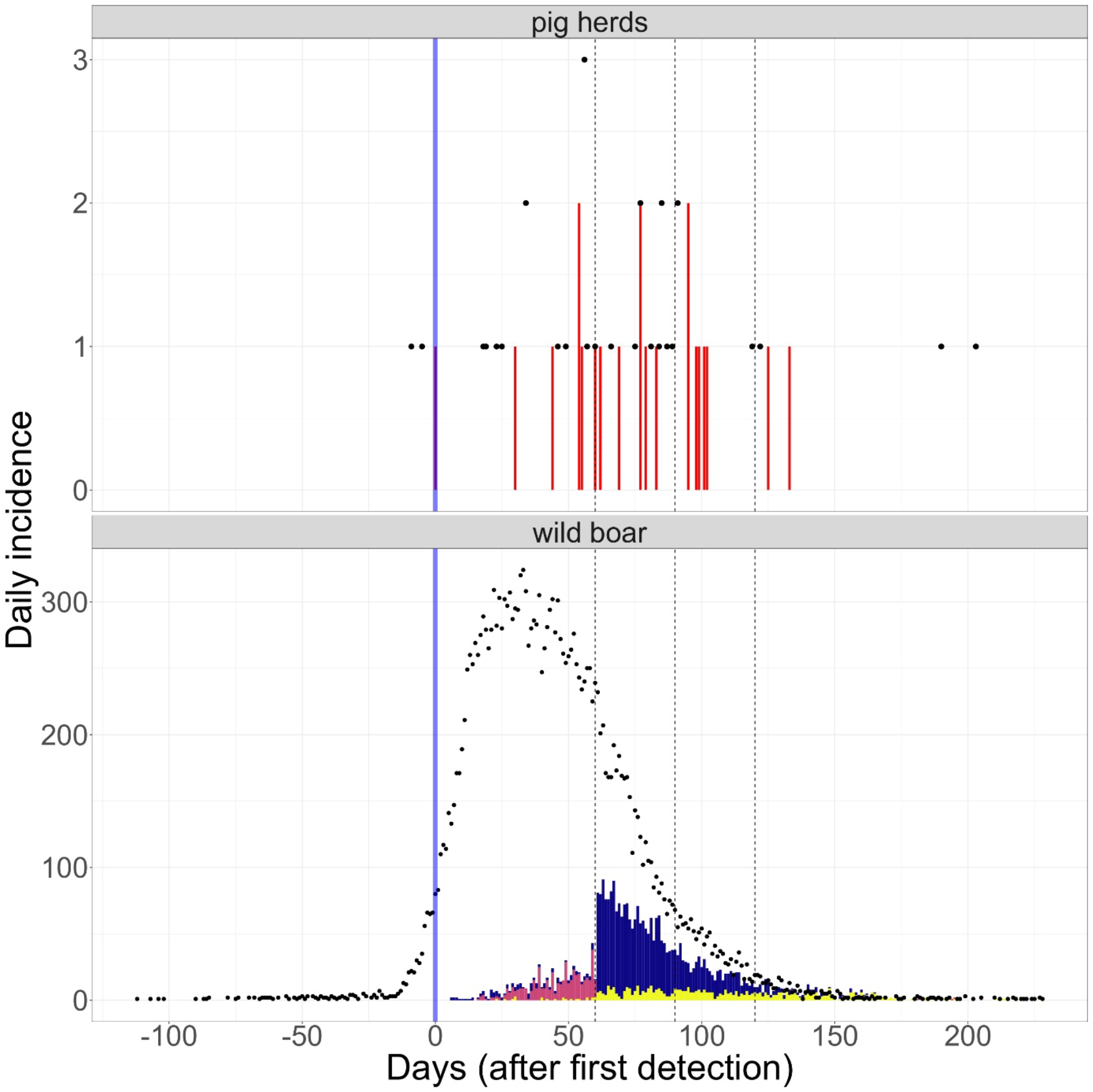
Total (dots) vs. detected (bars) number of new cases each day in pig farms (top) and wild boar (bottom) over time in the synthetic data provided to the ASF-Challenge teams. The blue vertical line represents the date of the first case detection. The cause of ASF detection is shown for wild boar: tests on hunted wild boar (blue) and infected carcasses found either through passive surveillance (yellow) or through active search (red). See supplementary information (SI8 and Movie S1) for a dynamic view of the synthetic epidemic.

The full datasets related to hunting bags, pig farms, live movements and ASF surveillance outputs for the three phases are available as supplementary material (see link to public repository in SI1).

#### 3.2.2 Situation reports

With each of the three data releases, narrative situation reports were provided to the teams. These reports aimed to contextualize the development of the epidemic during the phase that just passed and summarize the different control alternatives. They usually included:

- a description of the interventions put in place since the beginning of the phase;
- a narrative and qualitative summary of the epidemiological situation since the beginning of the phase;
- a description of what was expected from the modelling teams (see below).

A link to the three situation reports, released on days 50, 80 and 110 of the observed epidemic, is included in the supplementary information (SI7).

#### 3.2.3 Participant missions

At the end of phase 1, i.e. 80 days after the first detection, teams were asked to 1) predict the number and location of wild boar cases and outbreaks in farms that should be expected during the following four weeks; 2) predict the effectiveness of fencing the infected zone (the precise location of the fence was provided to the teams); and 3) advise on whether hunting pressure should be increased in the fenced area.

At the end of phase 2, i.e. 80 days after the first case detection, teams were asked to 1) update their predictions for the effectiveness of the fences implemented with or without the increased hunting pressure, now including a buffer zone of 15 km outside the fences; and 2) predict the effectiveness of five alternative control options, which were:

- Culling of all pigs in farms located in a protection zone;
- Increasing the size of the active search area around infected wild boar carcasses found outside the fenced/buffer areas (from 1 km to 2 km);
- Culling of all pigs in farms located within a 3 km radius of positive wild boar carcasses;
- Increasing the size of the surveillance zone (from 10 km to 15 km, but maintaining the surveillance period of 30 days);
- Culling of all pigs in farms that traded pigs with an infected farm less than three weeks before detection.

Finally, at the end of phase 3, i.e. 110 days after the first case detection, teams were asked to 1) update their predictions for the effectiveness of the fences implemented with or without the increased hunting pressure; 2) estimate the likelihood that the epidemic would fade out in the coming four months given the new control measures implemented; and 3) flag any long-term risk (i.e., second wave, risk of persistence in wild boar, etc.) and advise on how to mitigate this risk.

At the end of each phase, teams had six weeks to provide their model outputs and address the questions.

### 3.3 Expected predictions

After the end of the third phase, the forecasts and recommendations provided by the teams were analysed both qualitatively and quantitatively, and compared to synthetic data but also to what would have happened without additional control measures (Ezanno et al., submitted). Indeed, as measures implemented varied across phases of the ASF-Challenge, data provided for phase X+1 cannot be used to assess predictions from the teams based on data in phase X. Figure 6 shows the synthetic data provided to the ASF-Challenge teams together with the continuation of the prediction if nothing was changed in the next phase. This highlighted the impact of fences and increased hunting, which led to a doubling of the number of cases detected in wild boar (Fig. 6: phase 1). In contrast, the alternative measures tested with the model had a very low impact. While we kept the most effective one (preventive culling of pig in farms located within 3 km of any positive wild boar), the predictions with and without this complementary measure were very similar (Fig. 6: phase 2).

**Figure 6.**
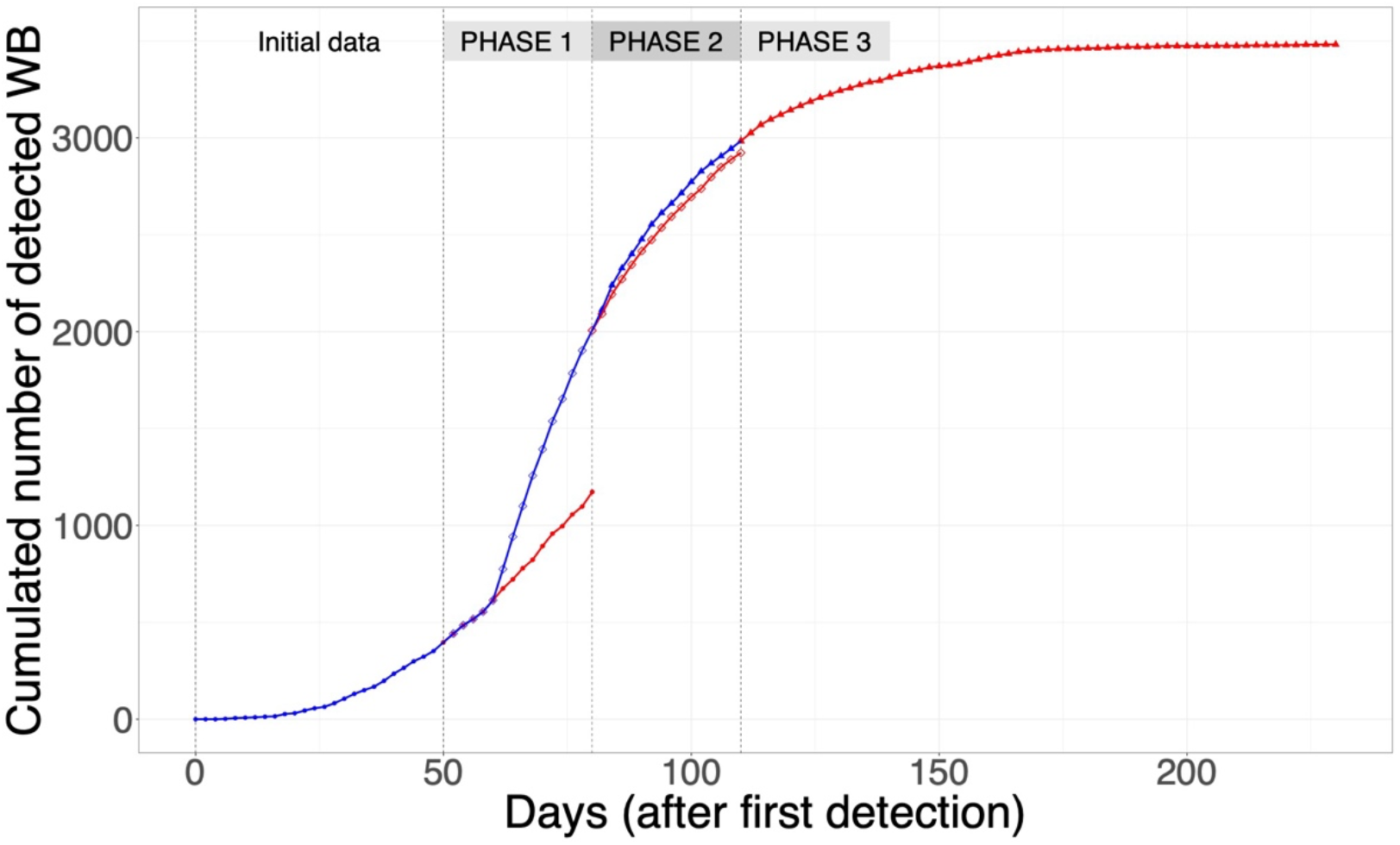
Synthetic data for the cumulative number of detected wild boar as provided to the ASF-Challenge teams (blue) and expected predictions with identical control conditions as the previous phase (red). Vertical dotted lines show the phase limits.

The very low number of infected wild boar detected 230 days after the first case detection (Fig. 5) could be seen as an indication of a highly probable epidemic fade-out. However, the distribution of the number of infected carcasses of wild boar (Fig. 7) at this same date clearly showed that the probability of extinction was very low.

**Figure 7.**
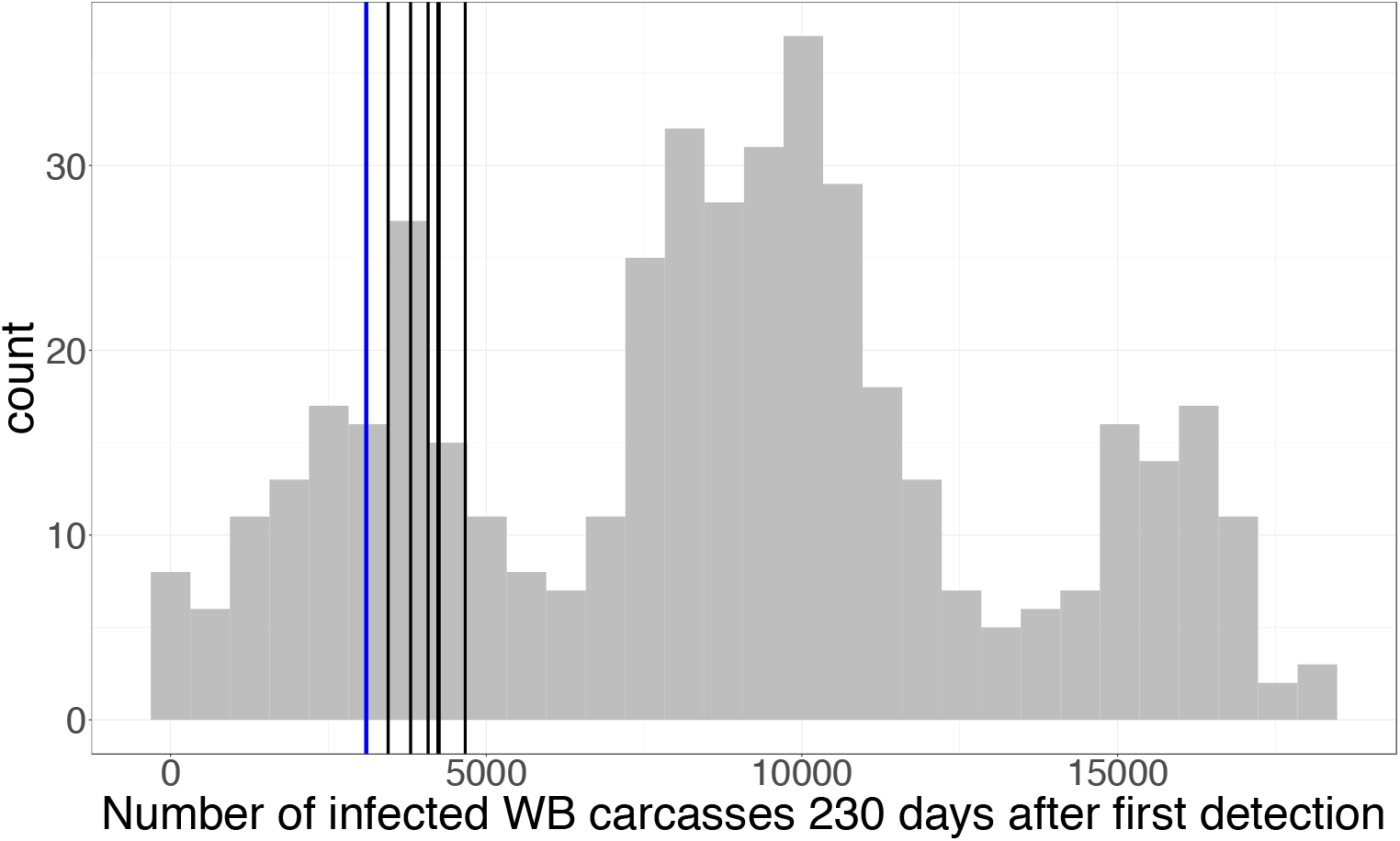
Distribution of the number of infected wild boar carcasses on the island 230 days after the first case detection over the 451 stochastic model repetitions. Vertical lines: in blue, repetition selected for the ASF-Challenge; in black, the 6 other repetitions meeting the five selection criteria.

## 4. Discussion

Several modelling challenges have been organized over the past 10 years on infectious diseases of public health concern, such as Ebola, influenza, dengue and chikungunya. They have been described as a unique framework allowing the development and evaluation of modelling approaches and forecasting methodologies that would not be possible in the context of retrospective analyses of historical epidemics (Ajelli et al. 2018). In the context of infectious livestock diseases, model comparisons efforts have been conducted, for instance for the foot-and-mouth disease (Probert et al., 2016; Webb et al., 2017), but no modelling challenges had been organized so far. This gap is particularly problematic since livestock populations and their pathogens have particular features that likely require specific modelling approaches and preparedness (Brooks-Pollock et al., 2015; Ezanno et al., 2020). This ASF modelling challenge was meant to fill this gap by allowing modelling teams to organize, get to know each other, develop new forecasting methodologies that could be deployed to investigate ASF epidemics, and compare their modelling approaches to that of the other teams.

For this first challenge, we focused on an ASF-like epidemic due to the global risk posed by this disease to the pork industry and the important role played by the wildlife reservoir in the epidemiology of the virus (Dixon et al., 2020). However, we did not aim to reproduce a “true” ASF epidemic since there seems to be almost as many ASF epidemic patterns as there are contextual situations (Sauter-Louis et al., 2021). In addition, in the current state of observed epidemics in Europe, fitting a transmission model on real data could be impeded by large reporting biases. Biological knowledge on ASF transmission dynamics must be strengthened before it is possible to faithfully reproduce what a real ASF outbreak would look like. Thus, we generated an epidemic that “looked like” an ASF epidemic by considering the relevant populations (domestic pigs and wild boar) and appropriate transmission processes within and between the two populations. Similarly, we have chosen to model between-farm animal movements in a fairly simple way, as we mainly focused on an intensive pig production system. Adapting this model for extensive or less structured systems with more backyard farms would certainly require more accurate patterns (Relun et al., 2017). We acknowledge that the calibrated model that was used to generate the synthetic data is only one of a virtually infinite set of models that could have generated ASF-like epidemic trajectories. In our simulated environment, the virus was detected much more frequently in wild boar than in domestic pig farms, which is consistent with several observed epidemiological situations such as in the Baltic States, Poland and the Republic of Korea (Sauter-Louis et al., 2021). Yet while it is important that the contextual situation represents a credible scenario inspired by real situations, there is no need to aim for a “perfectly calibrated” epidemic that likely does not exist.

To make the modelling challenge interesting and useful, the model that generated the synthetic data necessarily needed to be more detailed than the different models that were to be used to reproduce the data and make predictions. Since the challenge teams had not yet been recruited when the model was developed, we had no preconceived idea of the modelling approaches the different teams would use. To generate the synthetic data, we therefore developed a highly detailed stochastic individual-based transmission model with 93 parameters and seven transmission processes that very likely would be irrelevant for estimating transmission parameters and making useful predictions. The consequences of this complexity were that 1) the data generation and database preparation were computationally intensive; and 2) it was extremely time-consuming to calibrate the model in order to generate “ASF-like” epidemic trajectories that met the pre-defined selection criteria. In particular, the lack of data with regards to wild boar population dynamics and mobility patterns leaves no room for contrasting assumptions, and makes it difficult to anticipate how to account for an impact of fences on virus spread. The interface between livestock and wildlife, while crucial for better understanding multi-host diseases, is still poorly observed (Vicente et al. 2021).

Using synthetic data for a modelling challenge also renders it possible to control the “fog of war”, i.e. to know precisely the extent to which the detection data provided to modelling teams mirrors the infection data in both host populations. Here, we did not attempt to generate several scenarios for this “fog of war”, as done in the Ebola forecasting challenge (Viboud et al., 2018), since the objective was not to assess how fog of war could impact model predictions. Instead, we decided to create only one scenario that aimed to introduce sufficient uncertainty to mimic a realistic situation. A few pig farms were not initially in the farm database. Detection was far from perfect in both host populations, especially at the start of the epidemic and in wild boar.

## Author contribution

SP: Conceptualization; Formal analysis; Investigation; Methodology; Software; Supervision; Validation; Visualization; Writing the original draft

TV: Conceptualization; Data curation; Formal analysis; Investigation; Methodology; Supervision; Validation; Visualization; Writing the original draft

MM: Data curation; Formal analysis; Methodology; Investigation; Visualization; Writing - review & editing

SB: Formal analysis; Investigation; Methodology; Visualization; Writing - review & editing

PE: Conceptualization; Formal analysis; Funding acquisition; Investigation; Methodology; Project administration; Resources; Software; Supervision; Validation; Visualization; Writing the original draft

## Acknowledgements

C. Belloc & C. Peroz (Oniris), E. Gilot-Fromont (VetAgroSup), E. Baubet & E. Marboutin (OFB) are all thanked for their expertise on ASF, pig farm management, and wild boar population dynamics, which greatly helped us to define the model and associated scenarios.

We are most grateful to the Bioinformatics Core Facility of Nantes BiRD, member of Biogenouest, Institut Français de Bioinformatique (IFB) (ANR-11-INBS-0013), and to the INRAE MIGALE bioinformatics facility (MIGALE, INRAE, 2020. Migale bioinformatics Facility, doi: 10.15454/1.5572390655343293E12), for the use of their computing resources.

