## Supplementary Information for "The African swine fever modelling challenge: objectives, model description and synthetic data generation"

### 1 General information on the model

The model used to generate synthetic epidemiological data for the ASF Challenge was implemented using the EMULSION platform, version 1.1rc5, which requires Python (≥ 3.6) with library numpy (≥ 1.18) and can be installed through the 'pip' command. Details regarding principles, installation and usage of EMULSION are provided on the platform website: https://sourcesup.renater.fr/www/emulsion-public. Model files (ppa.py, ppa.yaml) and all the data and scripts required to run the model are available in the following public GIT repository: https://forgemia.inra.fr/spicault/asf-challenge-2020.

As the model is intended to represent a large area (about 500x500 km$^2$, i.e. about 1/4 of France), with more than 500,000 wild boar as spatially explicit individuals, fully susceptible epidemiological units (pig herds and wild boar) were represented only through vectors and matrix to spare computation time and reduce memory load. When becoming infected, each epidemiological unit was represented by an agent to provide the convenient detail level in managing the infectious process and control measures.

### 2 Between-unit dynamics

**Table S1.** Parameterization of the exponential transmission kernels between epidemiological units, depending on their type and status

| | | **Target of the force of infection** | |
|---|---|---|---|
| | | Alive wild boar | Pig farm |
| **Infectious source** | Alive wild boar | $\alpha_{bb}$ (direct contact): 2 km | $\alpha_{static}$ (outdoor only): 1 km |
| | Infectious wild boar carcass | $\alpha_{static}$ (direct contact) | *no transmission (no contact)* |
| | Pig farm | $\alpha_{static}$ (outdoor only) | $\alpha_{op}$ (other pathways): 1 km |

**Table S2.** Forces of infection experienced by each epidemiological unit, from other epidemiological units depending on their distance, type and status.

| | | **Target of the force of infection** | |
|---|---|---|---|
| | | Alive wild boar *w* | Pig farm *p* |
| **Infectious source** | Alive wild boar (*w'*) | $\beta_{bb} \times K_{\alpha_{bb}}(d(w,w')) \times \inf(w')$ | $\beta_{bp} \times K_{\alpha_{static}}(d(p,w')) \times inf(w') \times out(p) \times biorisk(p)$ |
| | Infectious wild boar carcass (*c*) | $\beta_{bb} \times K_{\alpha_{static}}(d(w,c)) \times \inf(c)$ | *no transmission (no contact)* |
| | Pig farm (*p'*) | $\beta_{bp} \times K_{\alpha_{static}}(d(w,p')) \times prev(p') \times out(p') \times biorisk(p')$ | $\beta_{bp} \times K_{\alpha_{static}}(d(p,p')) \times prev(p') \times biorisk(p') \times biorisk(p)$ |

Where:

- d(*a*, *b*) denotes the distance between epidemiological units *a* and *b*

- $K_\alpha(d) = e^{-\left(\frac{d}{\alpha}\right)^2}$ is an exponential transmission kernel, used with parameters $\alpha_{bb} = 2$ km, $\alpha_{static} = 1$ km, $\alpha_{op} = 1$ km depending on the transmission pathways (in practice, $K_\alpha$(d) was truncated to 0 for values below $10^{-9}$)
- inf($u$) denotes the contribution of epidemiological unit $u$ to virus transmission: 1 for fully infectious individuals (I, C), 0.5 for exposed (E)
- $prev(p) = \frac{0.5 \times E + I + C}{S + E + I}$ (where S, E, I, C denote the number of domestic pigs in the corresponding health states in farm $p$) is an equivalent of the classical prevalence accounting for the partial infectiousness of exposed animals
- out($p$) is 1 if pig farm $p$ provides outdoor access, 0 otherwise
- $biorisk(p) = (1 - biosec(p)) \times risk\_reduction$ where biosec($p$) is the biosecurity level of farm $p$ (1: perfect, 0: none) and *risk_reduction* a risk reduction factor to account for enhanced vigilance (1 in general, 0.5 in protection/surveillance zones or for traced herds)
- $\beta_{bb}$, $\beta_{bp}$, $\beta_{op}$ denotes, respectively, the transmission rates between wild boar, between wild boar and domestic pigs, and between pig farms (all values were set to 0.008)

#### 3 Detection

**Table S3.** Probabilities to detect infected domestic pigs depending on the nature of the farm and surveillance context. Column "Alert" applies to farms within a protection or surveillance zone, or identified at risk due to previous trade contacts with an infected farm (with trade ban and reinforced biosecurity).

| **Detection probability** (per infectious animal per day) | | **Epidemiological context** | | |
|---|---|---|---|---|
| | | Before 1st case | After 1st case | Alert |
| **Farm and animal type** | Commercial, alive | 0.01 | 0.02 | 0.1 |
| | Commercial, at death | 0.04 | 0.08 | 0.4 |
| | Backyard, alive | 0 | 0.005 | 0.025 |
| | Backyard, at death | 0 | 0.02 | 0.1 |

#### 4 Alternative control measures

The model was designed to make it possible to implement several additional control measures, either separately or in combination, and to trigger them at any moment in the model. In addition to those described in the paper, we tested the following actions:

1. Wider active search: when infected wild boar carcasses were found, other infected carcasses were searched within a 2 km-radius (instead of 1 km)
2. Wider surveillance zone: the radius of new surveillance zones was set to 15 km (instead of 10 km)
3. Cull in protection zone: when a protection zone is settled around a confirmed pig farm, all animals of the farms enclosed in the protection zone were preventively culled (and tested)

4. Cull traced herds: all animals of the farms in trade contact with an infected farm were preventively culled (and tested)

These measures were not retained in the setup of the challenge because they proved little efficacy in mitigating the spread of the virus in the simulated scenarios.

### 5 Model parameters

#### 5.1 Population dynamics

**Table S4.** Parameter values related to population dynamics.

| Symbol | Description | Value | Source |
|---|---|---|---|
| wildboar_death_rate | natural death rate of wild boar | 1/(365*5) $day^{-1}$ | Assumed |
| proportion_wildboars_hunted_default | proportion of the wild boar population hunted during the whole hunting period on the whole island | 0.5 | Assumed |
| hunting_start_date | delay between virus introduction and beginning of the hunting season | 42 days | Arbitrary choice |
| hunting_duration | duration of the hunting period on the island | 240 days | Assumed |

#### 5.2 Epidemiological processes

**Table S5.** Parameter values related to epidemiological processes.

| Symbol | Description | Value | Source |
|---|---|---|---|
| transmission_intra_commercial | transmission rate between pigs within a commercial pig farm | 0.4 $day^{-1}$ | Assumed |
| transmission_intra_backyard | transmission rate between pigs within a backyard pig farm | 0.6 $day^{-1}$ | (Halasa, 2016) based on experimental infections (Guinat, 2015), assuming a faster spread in unstructured herds |

| | | | |
|---|---|---|---|
| transmission_boar_pig ($\beta_{bp}$) | transmission rate between wild boar and pigs | 0.008 $day^{-1}$ | Assuming transmission rate is the same between species if contact occurs, contacts being modelled by kernels |
| transmission_boar_boar ($\beta_{bb}$) | transmission rate between wild boar | 0.008 $day^{-1}$ | Same |
| transmission_other_pathways ($\beta_{op}$) | transmission rate from farm to farm due to indirect contacts | 0.008 $day^{-1}$ | Assumed |
| wildboar_C_removal_rate | rate at which wild boar carcasses disappear in nature | 1/90 $day^{-1}$ | Assumed |
| external_risk_reduction | multiplicative factor for reducing indirect transmission within pig farms and between pigs and wild boar, due to improved biosecurity. 0: no external risk, 1: unchanged risk | 0.5 | Assumed |
| incubation | average duration in the incubating state (E) for pigs and wild boar | 7 days | Assumed |
| asf_death_rate | mortality rate in the infectious state (I) for pigs and wild boar | 1/7 $days^{-1}$ | Assumed |
| contrib_E_transmission | ratio of E shedding rate over I shedding rate | 0.5 | Assumed |

#### 5.3 Detection and control (regulatory measures)

**Table S6.** Parameter values related to detection of infected epidemiological units and regulatory control measures.

| Symbol | Description | Value | Source |
|---|---|---|---|
| pig_contact_window | duration during which all past trade contacts (in/out) between a farm and an infected farm are considered at risk (days) | 21 days | Regulatory measures |

| | | | |
|---|---|---|---|
| proba_detection_hunted_wildboars_default | default proportion of hunted wild boar tested once first case detected | 0.2 | Assumed |
| proba_boar_detection_carcass_primary | probability to find and detect each infected wild boar carcass each day as long as no primary case is found | $10^{-4}$ | Assumed |
| proba_boar_detection_carcass_secondary | probability to find detect infected wild boar carcasses when a first case has already been detected | $10^{-4}$ | Assumed (no impact of primary case on WB passive surveillance) |
| proba_detection_carcass_active_search | probability that a wild boar carcass (infected or not) is found when an active search is performed within a given area (probability for the whole search period) | 0.1 | Assumed |
| default_radius_boar_detection_carcass_active | default radius of the area for active carcass search around a case | 1 km | Assumed |
| delay_carcass_active_search | delay between the discovery of an infected wild boar carcass and the organization of an active wild boar carcass search in the corresponding area | 5 days | Assumed |
| duration_carcass_active_search | duration of the active search campaign to find wild boar carcasses | 5 days | Assumed |
| delay_confirmation | number of days needed between suspicion and confirmation in pig farms, pig removal being then immediate | 4 days | Assumed |
| delay_boar_removal | delay to remove wild boar carcasses found in nature | 0 days | Assuming that carcasses are immediately removed to be tested (before any test results) |
| repopulation_delay | delay after which a pig herd which was entirely removed after confirmation can be repopulated | 50 days | Regulatory measures |

| | | | |
|---|---|---|---|
| trade_suspicion _duration | duration during which herds in contact with confirmed cases are considered suspicious ("traced farms"), assumed equal to the duration of the protection zone | 40 days | Assumed |
| protection_radius | radius of the protection zone, centred on a confirmed case, within which all pig herds are considered suspicious | 3 km | Regulatory measures |
| protection_duration | duration during which herds within the protection zone are considered suspicious | 40 days | Regulatory measures |
| surveillance_radius | radius of the surveillance zone, centred on a confirmed case, within which all pig herds are considered suspicious | 10 km | Regulatory measures |
| surveillance_duration | duration during which herds within the surveillance zone are considered suspicious | 30 days | Regulatory measures |

#### 5.4 Alternative control measures

**Table S7.** Parameter values related to alternative control measures (all assumed).

| **Symbol** | **Description** | **Value** |
|---|---|---|
| radius_wider_active_search | enlarged radius of active search for wild boar carcasses | 2 km |
| delay_implementation | duration between the decision to perform a preventive culling and the actual culling | 3 days |
| radius_cull_near_infected _carcasses | radius around infected wild boar carcasses to implement preventive culling of pig herds | 3 km |
| delay_cull_near_infected _carcasses | delay between detection of primary case and the implementation of the preventive culling of pig herds near infected wild boar carcasses | 90 days |
| radius_wider_surveillance_area | enlarged radius for surveillance zone | 15 km |
| fences_efficacy | coefficient used to modify kernels when installing fences. Affected kernels are multiplied by (1-fences_efficacy), thus 1 is the highest efficacy (kernel becomes 0), 0 the lowest (kernel unchanged) | 0.99 |

| | | |
|---|---|---|
| delay_use_fences | delay between detection of primary case and full installation of fences | 60 days |
| proportion_wildboars _hunted_fences | proportion of the wild boar population hunted within fences between the decision to reduce population and the end of hunting period | 0.9 |
| buffer_area_size | Size of the buffer area around fences | 15 km |
| proportion_wildboars _hunted_buffer | proportion of the wild boar population hunted around fences between the decision to reduce population and the end of the special population reduction period | 0.9 |
| duration_hunting_within_buffer | duration (days) of increased hunting on the tiles surrounding fences | 61 days |
| proba_detection_hunted_wildboars _increased | proportion of hunted wild boar tested within fences and buffer | 1 |
| factor_carcass_detection _increased_hunting | factor which multiplies the probability to detect wild boar carcasses within/around fences when increased hunting is implemented | 5 |

### 6 Stochastic simulations and selected epidemic trajectory

**Table S8.** Selection criteria for choosing the epidemic trajectory used in the Challenge. Proportions are relative to the 451 repetitions kept after discarding repetitions without any detection or with detection of the primary case later than 200 days after virus introduction.

| Criterion | Rationale | Number (%) of repetitions |
|---|---|---|
| > 250 infected wild boar before detection of primary case | Disease sufficiently established in wildlife | 340 (75.4 %) |
| Primary detection in pig farm | Most probable 1st detection event because of low passive surveillance in wild boar before any 1st case | 153 (33.9 %) |
| < 500 wild boar infected outside the fenced area at installation | Possible effectiveness of fences | 230 (51 %) |
| > 250 wild boar infected 110 days after detection of primary case | Phase 3 of Challenge achievable | 418 (92.7 %) |
| < 30 infected wild boar 230 days after detection of primary case | Positive message at the end of the challenges | 91 (20.2%) |

#### 7 Situation reports

Situation reports regarding days 50, 80 and 110 after the detection of the first case were provided to the participants. They are available on the public git repository: https://forgemia.inra.fr/spicault/asf-challenge-2020.

#### 8 Dynamic view of the epidemics

File Movie_S1.mp4 provides an dynamic view of the epidemics spread (on the left), with infected wild boar in black and infected pig farms in blue (the first infected wild boar and the primary case which is a pig farm are identified through a star and a square, respectively), and of the temporal evolution of infection and detection (on the right) with the same legend as in Figure 5.
